# The mitochondrial DNA content can not predict the embryo viability

**DOI:** 10.1101/445940

**Authors:** M.S. Yong Qiu, M.S. Songchang Chen, Chen Dayang, M.S. Ping Liu, M.S. Jun Xia, B.S. Lin Yang, M.S. Zhe Song, M.S. Qianyu Shi, M.S. Lin Xie, M.S. Zhu Zhu, Du Ye, Hui Jiang, Jian Wang, Huanming Yang, Chenming Xu, Fang Chen

## Abstract

**Objective:** To investigate whether the mitochondrial DNA content could predict the embryo viability

**Design:** Retrospective analysis.

**Setting:** Reproductive genetics laboratory

**Patient(s):** A total of 421 biopsied samples obtained from 129 patients

**Intervention(s):** Embryo biopsies samples underwent whole genome amplification (WGA) and were tested by next generation sequencing (NGS) and array Comparative Genomic Hybridization (aCGH), 30 samples were selected randomly to undergo quantitative real-time polymerase chain reaction (qPCR).

**Main Outcome Measure(s):** Those embryos which obtained the consistent chromosome status determined both aCGH and NGS platform were further classified. We investigated the relationship of mtDNA content with several factors including female patient age, embryo morphology, chromosome status, and live birth rate of both blastocysts and blastomeres.

**Result(s):** A total of 386 (110 blastomeres and 276 blastocysts) out of 399 embryos showed consistent chromosome status outcome. We found no statistically difference was observed in aneuploid and euploid blastocysts (p=0.14), the same phenomenon was observed in aneuploid and euploid blastomeres (p=0.89). Similarly, the mtDNA content was independent of female patient age, embryo morphology and live birth rate.

**Conclusion(s):** The mtDNA content did not provide a reliable prediction of the viability of blastocysts to initiate a pregnancy.

## Introduction

Identification and development of strategies for increasing implantation and pregnancy rates are an important scientific issue in the field of assisted reproduction (1). With the help of elegant grading systems combined with preimplantation genetic screening (PGS) approaches based on determination of embryo DNA content, a lot of clinical trials suggested that euploid embryo showed higher pregnancy and implantation rates (2, 3). However, there was still a significant percentage of morphologically and chromosomally normal embryos failed to implant, suggesting there must be other factors affecting embryo viability(4, 5).

In the process of embryo development, most cellular energy is derived from the mitochondria (6–9). Therefore, adequate energy maybe contributed to embryo viability (5). Until now, several groups have demonstrated that the mitochondria DNA (mtDNA) influences the potential of fertilization (10, 11). Fragouli et al. suggested mtDNA levels could provide an independent measure of embryonic implantation potential (12). Ravichandran et al. assessed that mtDNA quantification as a tool for embryo viability using 1505 embryos from 35 IVF clinics. (13). Recently, the mtDNA content was further indicated to be negatively related to embryo viability and proposed to be used as a biomarker in the prediction of implanation outcome (4, 12, 14, 15). But these results were still under debate (16–18). Victor et al. reported a modified method that demonstrated mtDNA lacked of predictive value (17). Three independent studies presented that mtDNA content failed to establish predictive power of implantation at the conference of Preimplantation Genetic Diagnosis International Society (PGDIS) (19–21). Moreover, Treff et al. demonstrated that levels of mtNDA didn’t predict the reproductive potential in the similar gold standard : double embryo transfer (DET)(18). What is the cause of this inconsistency? That maybe caused by technical defiencies or original data. Firstly, Fragouli et al. performed mtDNA content quantification using a multi-copy sequence (Alu) as normalization (12). It is worth noting that many multi-copy sequences (such as Alu) which comprise self-transposable elements, that may cause variability among the human [28, 29]. Secondly, Reprogenetics group’s data showed mtDNA content lacked of predict power in 17 out of 35 clinics that demonstrated anything linking mtDNA content and implantation was not a universal biological phenomenon (16). not scientific for analyzing the mtDNA content in embryo which came from many IVF clinics together. Thirdly, the previous studies just used next generation sequencing (NGS) or array Comparative Genomic Hybridization (aCGH) to determine each embryo chromosome status(12, 13, 18), it is worthy to note that the consistency of the results based on NGS and aCGH platform is not perfectly 100% (22–24). That also could cause some deviations when do classification based on embryo chromosome status.

In our study, we quantified mtDNA content using single copy sequence (GAPDH) as normalization to avoid the variability in human population. Besides, all of the embryos in our study were from one IVF clinics. Additionally, we used both NGS or aCGH to determine the embryo chromosome status accurately. Based on above strategy, we investigated the relationship between mtDNA content and several factors including chromosome status, embryo morphology, female patient age, live birth rate. Our results demonstrated that the mtDNA content was independent of embryo chromosome status, embryo morphology and female patient age. Moreover, no statistically significant difference was observed in the mtDNA content between embryos with and without producing live birth. Based on these results, we concluded that the mtDNA content of blastocysts did not predict the embryo viability.

## Material and Methods

### Samples Collection and Ethical approval

A total of 421 biopsied samples were collected for assisted reproduction treatment procedures in The International Peace Maternity & Child Health Hospital of China welfare institute (IPMCH). And obtained the approval of the Institutional Review Board of both BGI-Shenzhen and IPMCH. All patients have participated in this trial on an informed-choice basis and have signed a consent form for the purpose of this test.

### Embryo Biopsy and morphological criteria

Embryo biopsies were performed using laser-assisted biopsy at the cleavage or blastocyst stage in all cases, as previously described (25, 26). Embryo grade is determined according to elegant grading systems based on embryo cleavage rate and morphology (27).

### WGA and Array-Comparative Genomic Hybridization

WGA was performed using the PicoPLEX™ WGA Kit(Rubicon Genomics, Ann Arbor, MI, US) according to the manufacturer’s protocol (28), the same WGA kit to previous similar study(12). The aCGH was carried out using 24 Sure Cytochip V3 microarrays (Illumina Ltd., Cambridge, UK). The protocol used was as described in Fragouli et al. (29).

### Library preparation and sequencing

We prepared libraries starting with 300 ng totals amplified DNA for each sample. Firstly, we fragment DNA using NEBNext^®^ dsDNA Fragmentase^®^ (NEB, Ipswich, MA, USA). Then the fragment DNA was purified using beads. The following steps were conducted according to the BGISEQ-500 protocol to give the final library and sequencing (30, 31).

### Bioinformatics Analysis of Chromosomal Abnormality

The 50bp reads were subsequently aligned to the human genome hg19 (GRCh37, UCSC release hg19) using the SOAP2 (32), allowing a maximum of two base pair mismatches. After alignment, we picked out the unique non-duplication mapping reads from the SOAP file. The genome sequence was divided into unequal length windows according to an optimized dynamic observation windows selection strategy and the number of reads per window was calculated. For CNV detection, we referred to an algorithm developed in BGI-Shenzhen (33). In brief, the number of reads per window was normalized by the average reads number in all autosome windows. The normalized results were defined as copy ratio for each window. The average copy ratio of the whole Y chromosome was used for sex determination, with 0.001 as a threshold. For aneuploidy detection, the mean copy ratio of whole chromosome was used to determine the aneuploidy status.

### Calculation the mtDNA content using NGS data

Relative amounts of mtDNA and nuclear DNA were determined by NGS. The mitochondrial reads were extracted from the SOAP files, which uniquely aligned the mitochondrial reference sequence and removed the duplication reads. The whole effective genome reads were the unique non-duplication mapping reads of nuclear DNA. The mtDNA content was calculated based on NGS data refer to Victor et al.(17). The ratio of mitochondrial reads is multiplied by the correction factor F_NGS_ as per equation.

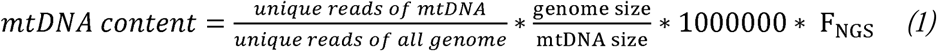

F_NGS_ take into embryo gender and ploidy to correctly normalize for the ratio of mitochondrial reads.

### Determine the mtDNA Content using qPCR

The mtDNA content determination was performed using the multiple QPCR TaqMan assay. A probe (5’-FAM-GCGTTCAAGCTCAACACCCACTACC-BHQ1-3’, Life Technologies, UK) was designed to target mtDNA region. A second probe (5’-JOE-GGCTGGGGCCAGAGACTGGC-BHQ1-3’, Life Technologies, UK) targeting GAPDH sequence representation of nuclear DNA as the normalization. Two repeats were set up for both the mtDNA and GAPDH sequences. The reaction and condition were performed according to the previous study (12).

## Results

### Data production

A total of 18.65 ± 5.46 million reads were obtained for each biopsied sample on BGISEQ-500, covering 7.47 ± 1.70% of the whole human genome. 82.51 ± 2.56% reads were mapped to the reference, while 64.45 ± 2.26% reads were uniquely mapped after removing duplicated reads, with duplication rate of 5.82 ± 1.18%. The GC content was 42.71 ± 1.20% in accordance with that of the human reference genome. We also calculated the data mapped to mitochondria DNA. After removing duplicated reads, average 2454.31 ± 1596.56 reads uniquely mapped to mitochondria DNA, covering 55.36% ± 14.19% of the whole mitochondrial DNA and the duplication rate is high up to 84.99% ± 7.62%.

### MtDNA content in groups classified on embryo chromosome status

We performed both NGS and aCGH to determine the embryo chromosome status and 386 (110 blastomeres and 276 blastocysts) out of 399 embryos showed consistent chromosome status by these two platforms (Fig 1) and were further classified into euploid or aneuploid chromosome constitution. Then we calculated the mtDNA content in 140 aneuploid (73 blastocysts and 67 blastomere) and 246 euploid (203 blastocysts and 43 blastomere) embryos, respectively (Supplementary Table I). Simultaneously, to verify the accuracy of mtDNA quantitation based on NGS data, a cross-platform of real-time polymerase chain reaction (qPCR) was used to test 30 samples randomly. We found the coefficient was 0.61 using Pearson’s chi-squared test, which suggested that the same trend was observed in mtDNA content in these two measurement methods (Supplementary Fig. 1).

As results, we found no statistically difference was observed in aneuploid and euploid blastocysts (p=0.14), the same phenomenon was observed in aneuploid and euploid blastomeres (p=0.89) (Figure 2A). That suggested the mtDNA content was independent of embryo chromosome status.

**Figure 1.**
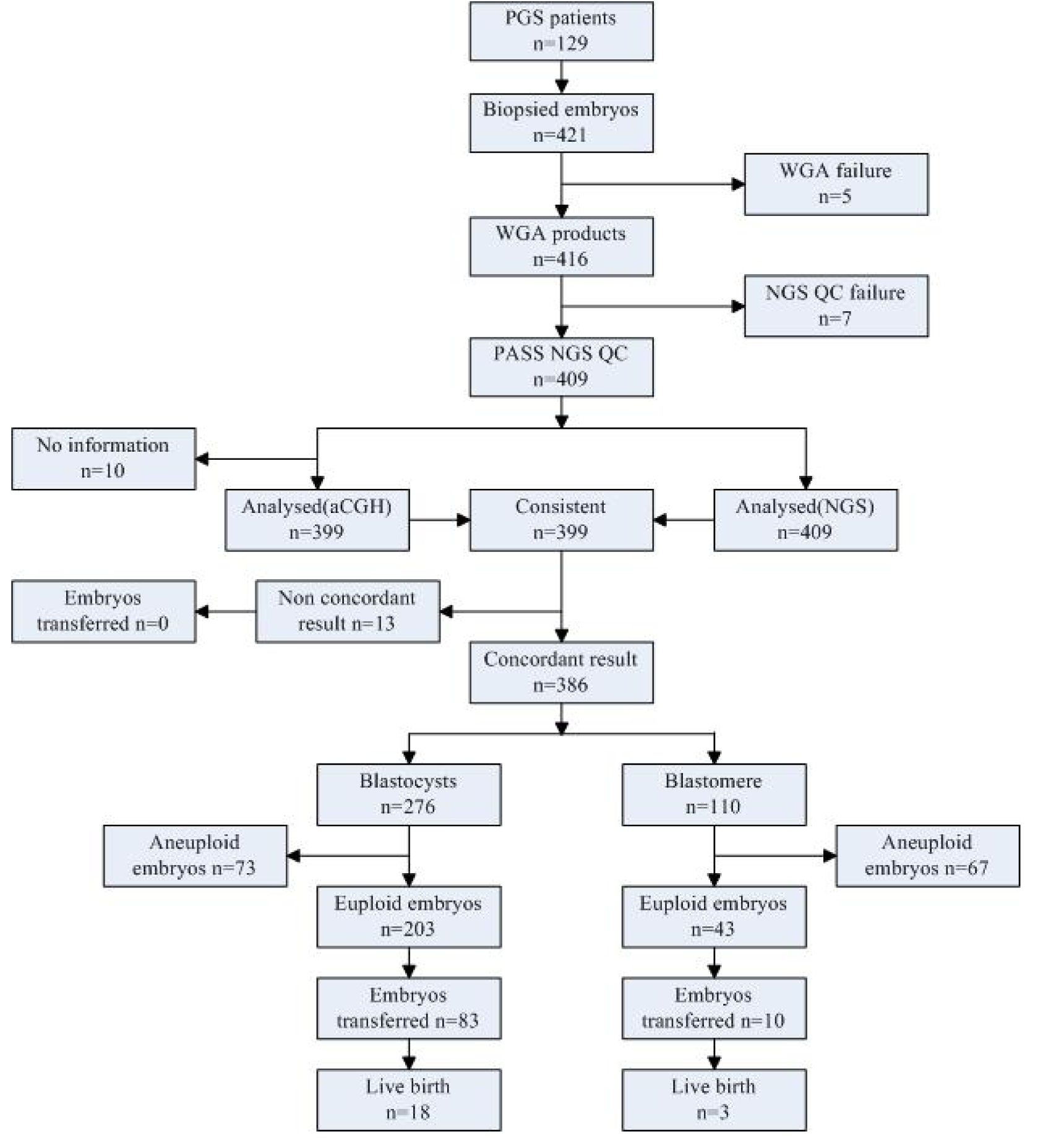
The workflow of this study. PGS, preimplantation genetic screening; WGA, whole genome amplification; QC, quality control; aCGH, array comparative genomic hybridization; NGS, next-generation sequencing.

### The mtDNA content was independent of female patient age and embryo morphology

To explore the relationship between the mtDNA content and female patient age, we performed Loess analysis on the mtDNA content in 203 euploidy blastocysts (patient age ranged from 24 to 40 years) and 43 euploidy blastomeres (patient age ranged from 25 to 41 years). The regression line showed that there was no linear correlation between mtDNA content and female patient age (in blastomeres, p = 0.46, r = -0.12, in blastocysts, p = 0.9, r = 0.0092 Pearson correlation) (Fig 2B).

Embryo morphology is a key reference factor in selecting embryos for IVF transferring. It was therefore of interest to investigate the relationship between the mtDNA content and embryo grades determined mainly based on morphology. Owing to the small data of inner cell mass A(ICM_A), a total of 276 blastocysts trophoectoderm (TE) and 110 blastomeres were used for this evaluation (Supplementary Table 2). But no statistically significant difference was observed among trophoblast different grades (Fig 2C), the same conclusion was obtained among blastomere different grades (Fig 2D). Based on above results, that suggested the mtDNA content was independent of embryo morphology.

**Figure 2.**
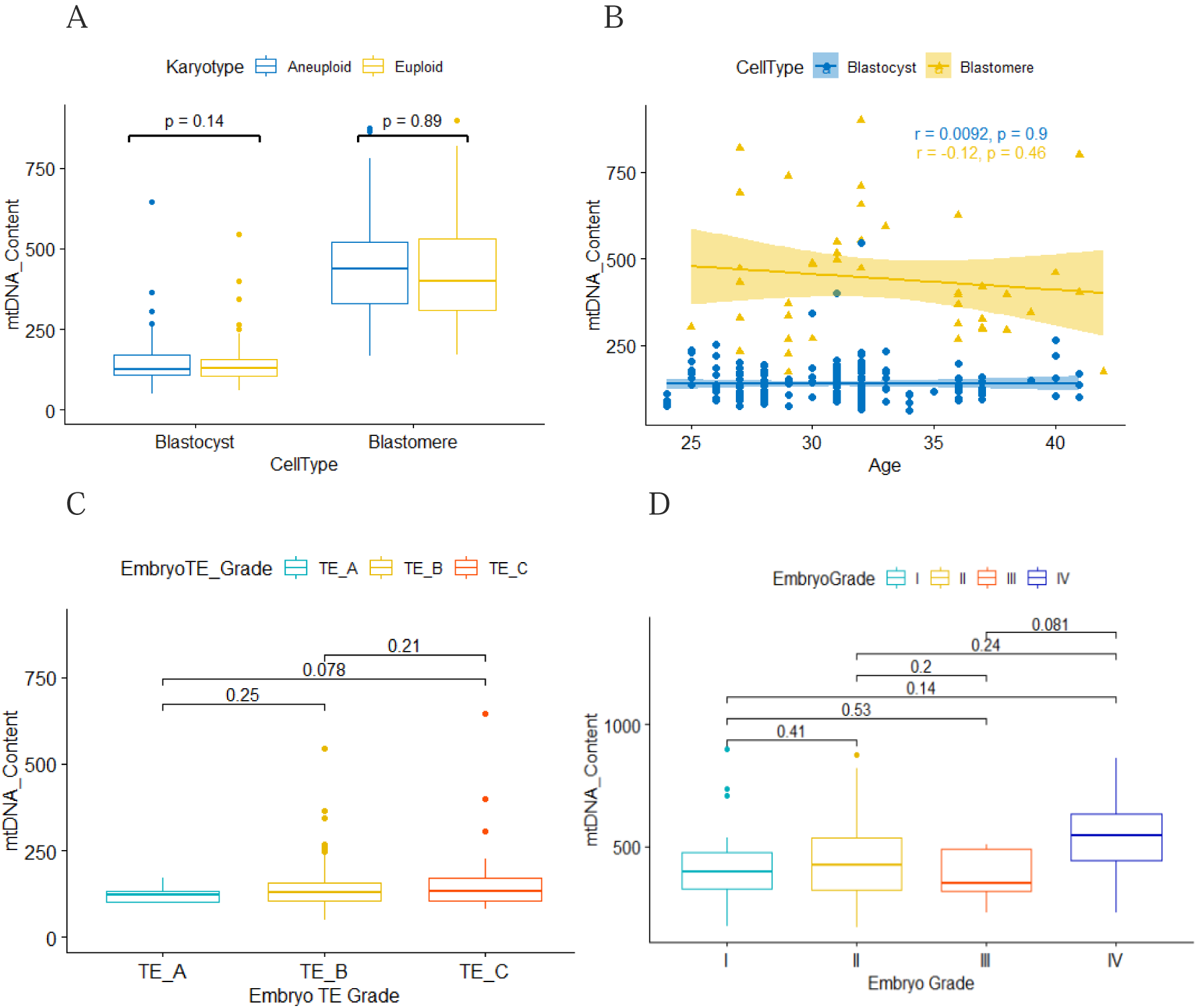
The mtDNA content in relation to embryo chromosome status, patient age and embryo grade. (A)The embryos were classified based on chromosome status. (B) The embryos were classified based on patient age. (C)The embryos were classified into trophectoderm (TE) grading and (D) blastomere grading, respectively. The y-axis represented the value of mtDNA content. The x-axis represented the embryo parameters.

### The mtDNA content couldn’t predict the live birth

In order to assess whether mtDNA content had an influence on the viability of an embryo, we compared the mtDNA content in three groups blastocysts which were classified based on implantation results. The three groups are “N” groups (no transferred embryos, n=120), “F” groups (embryos failed to produce live birth, n=65) and “S” groups (embryos produce live birth successfully, n=18) respectively. Among 83 transferred blastocysts, two blastocysts have no follow-up information, eighteen embryos produced live birth successfully.

Owing to only 3 blastomere produced live birth successfully, only the mtDNA content in blastocysts was investigated. Data showed that no statistically significant difference was observed among these three groups (Figure 3). Although the live birth rate was 18 out of 81, we still concluded that the mtDNA content did not provide a reliable prediction of a blastocyst to produce a live birth, but it requires more data to support.

**Figure 3.**
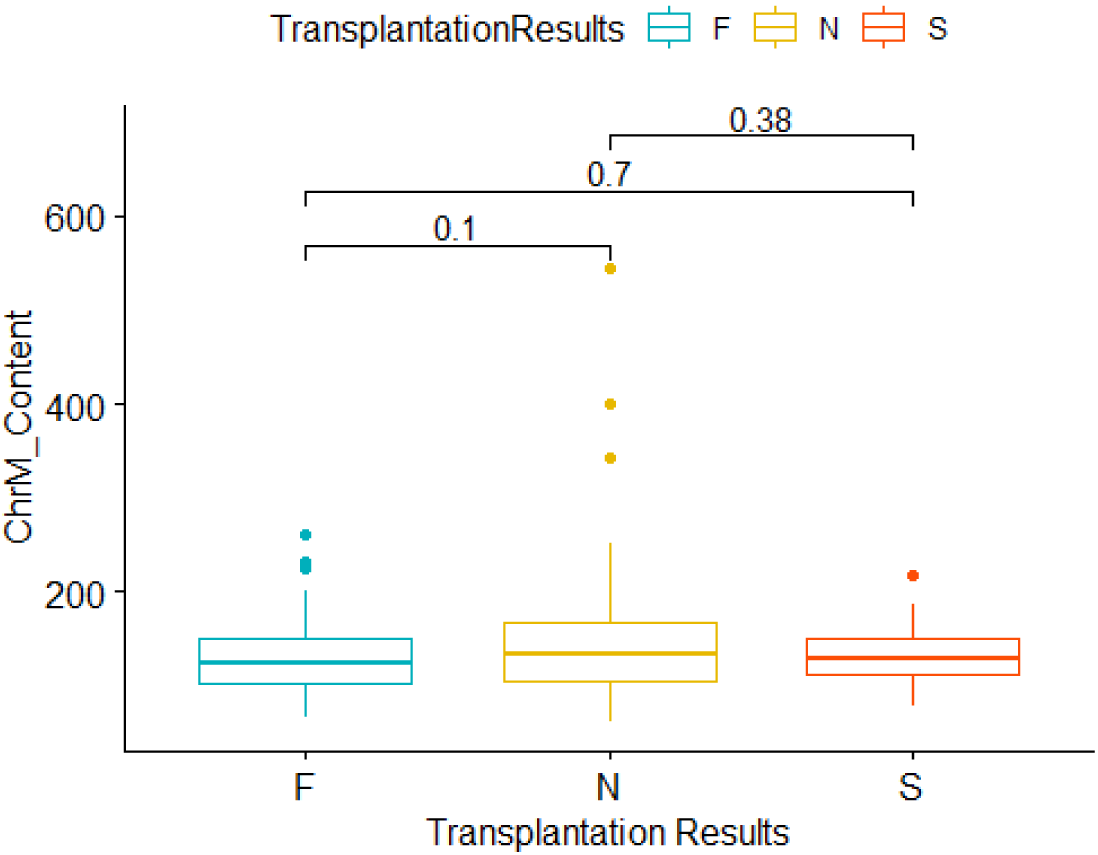
The mtDNA content in relation to live birth. N represented no transferred embryos. F represented embryos transferred but failed to produce a live birth. S represented embryos produce live birth successfully. The y-axis represented the value of mtDNA content. The x-axis represented the embryo parameters. Additionally, in order to reduce confused factors in our study, we investigated 18 patients who underwent more than one transfer from the same embryo cohort and compared the mtDNA content between the blastocyst that produced live birth and those did not (Supplementary Fig. 2). This analysis showed that mtDNA content didn’t predict the viability of an embryo effectively.

## Discussion

Our study finds no correlation of mtDNA content with embryo chromosome status, female patient age, embryo grade and live birth rate, that suggested mtDNA content couldn’t predict the embryo viability.

Whether mitochondrial DNA (mtDNA) copy number could serve as a biomarker for embryo viability was a hot debate. In this debate, several groups believed that mtDNA content could be a tool for embryos choosing(14, 15). But other groups demonstrated that mtDNA content didn’t provide a value predictor(17, 18). Why mtDNA content could be a predictor in some center but not in others? That maybe caused by technical defiencies or original data. Firstly, the quantifying mtDNA content accurately was a key step in this debate. The majority researchers used qPCR or NGS platform to quantify the mtDNA content. For qPCR, it mainly relies on a ratio of two values (mtDNA and nuclear DNA). In this study, we choose a single copy loci GAPDH represent nuclear DNA instead of multi-copy sequences (Alu) which comprise self-transposable elements could cause fluctuation in human population [28, 29]. Simultaneously, we use a cross-platform (qPCR and NGS) to validation the measurement accuracy. The regression line showed that there was linear correlation between mtDNA content using NGS platform and qPCR platform (p = 6e-04, r = 0.61, Pearson correlation) (Supplementary Fig. 1).

Secondly, Reprogenetics group’s data showed mtDNA content lacked of predict power in 17 out of 35 clinics that demonstrated anything linking mtDNA content and implantation was not a universal biological phenomenon (16). That suggested it was not scientific for analyzing the mtDNA content in embryo which came from many IVF clinics together. In our study, we just analyzed the embryos form one IVF clinics. Lastly, in the previous studies, only aCGH or NGS technology was used to determine the embryo chromosome status (4, 12, 18, 34). However, it is worthy to note that the consistency of the results based on NGS and aCGH platforms is not perfectly 100% (22–24). To settle this issue, in this study, we simultaneously used NGS and aCGH methods to determined the chromosome status, and only selected these embryos with consistent results to perform further analysis. That means the classification based on the chromosome status was much accurate. But there was still no statistically difference observed in aneuploidy and euploidy embryos.

In addition, we investigated 19 patients that had undergone more than one transfer from the same embryo cohort and compared mtDNA content between the embryos that produced a live birth and those that did not. Also in this study, mtDNA content was not a useful predictor of live birth (Supplementary Fig. 2).

Anyway, although there were some limits existed in our study, such as the fluculation in unique reads mapped to mtDNA in embryos and the small number of embryos producing live birth. Only 18 out of 81 embryos produced live birth successfully in our study, the birth rate was 22%, that was lower than expected. We still would like to conclude there was no link between mtDNA content and embryo viability.

## Acknowledgements

The authors would like to thank the medical staff in IVF clinics for their assistance with clinical outcome data collection. The authors are very grateful to the staff in MGI Tech Co.,Ltd. for their advice on sequencing platform.

